# Genomic Diversity in a Population of Spodoptera frugiperda Nucleopolyhedrovirus

**DOI:** 10.1101/2020.10.27.358317

**Authors:** Tomás Masson, María Laura Fabre, Matias Luis Pidre, José María Niz, Marcelo Facundo Berretta, Víctor Romanowski, María Leticia Ferrelli

## Abstract

Spodoptera frugiperda multiple nucleopolyhedrovirus (SfMNPV) represents a strong candidate to develop environmental-friendly pesticides against the fall armyworm (*Spodoptera frugiperda*), a widespread pest that poses a severe threat to different crops around the world. However, little is known regarding the genomic diversity present inside SfMNPV isolates and how it shapes the interactions between virus and host. Here, the genomic diversity present inside an isolate of SfMNPV was explored using high-throughput sequencing for the first time. We identified 704 intrahost single nucleotide variants, from which 184 are nonsynonymous mutations distributed among 82 different coding sequences. We detected several structural variants affecting SfMNPV genome, including two previously reported deletions inside the *egt* region. A comparative analysis between polymorphisms present in different SfMNPV isolates and our intraisolate diversity data suggests that coding regions with higher genetic diversity are associated with oral infectivity or unknown functions. In this context, through molecular evolution studies we provide evidence of diversifying selection acting on *sf29*, a putative collagenase which could contribute to the oral infectivity of SfMNPV. Overall, our results contribute to deepen our understanding of the coevolution between SfMNPV and the fall armyworm and will be useful to improve the applicability of this virus as a biological control agent.

**Highlights:**

- We characterized the genomic diversity within a population of SfMNPV.
- Coding regions with higher genetics diversity are associated with oral infectivity or unknown functions.
- Several structural variants contribute to the genomic diversity of SfMNPV.
- *Sf29*, a putative collagenase, shows signs of adaptive evolution.

## 1. Introduction

The fall armyworm, *Spodoptera frugiperda* (J.E. Smith, Lepidoptera: Noctuidae), is an invasive agricultural pest that attacks a variety of crops and is naturally distributed across the Americas. Recent reports have also described the presence of *S. frugiperda* in the African and Asian continents (Stokstad 2017; Silver 2019). Traditional control methods for this pest involve the use of chemical products or crops expressing Bt toxins (Bernardi et al. 2015). However, cases of field-evolved resistance to Bt toxins and environmental impact of chemical pesticides are forcing the development of novel control strategies (Harrison et al. 2019). Baculoviruses are insect-specific large DNA viruses (Harrison et al. 2018) with a narrow host range and high virulence, which make them potentially safe candidates to develop biopesticides (Haase, Sciocco-Cap, and Romanowski 2015). Specifically, a nucleopolyhedrovirus infecting *S. frugiperda* (SfMNPV) has been implemented for the biological control of fall armyworm populations (Bentivenha et al. 2018).

To date five SfMNPV complete genomes have been reported (Harrison, Puttler, and Popham 2008; Wolff et al. 2008; Simón et al. 2011; Simón, Palma, et al. 2012; Barrera et al. 2015), opening the door to comparative analysis that explores the relationship between virus genome evolution and virulence-associated mutations. Deletions and mixed genotypes are common in natural populations of SfMNPV and have been proposed as modulators of its speed of killing and potency (Clavijo et al. 2009; Simón, Williams, et al. 2012).

Genetic innovation is one of the main drivers of adaptation during virus-host coevolution processes (Obbard and Dudas 2014; Holmes 2011). For the case of the model baculovirus, *Autographa californica* multiple nucleopolyhedrovirus (AcMNPV), population polymorphism has proven to be a relevant source of functional diversity (Chateigner et al. 2015). Moreover, it has been proposed that escape from host suppression is driven by a subset of genes under recurrent positive selection (Hill and Unckless 2017). Despite the availability of consensus sequences, the genomic diversity present within SfMNPV isolates have remained uncharacterized. This lack of information limits our knowledge of the adaptation process of SfMNPV to the fall armyworm.

In this work, we address this issue through the characterization of the genome-wide diversity present inside an argentinian isolate of SfMNPV (ARG-M) by deep sequencing. Through the analysis of nonsynonymous variants, we identified genes related to either oral infection or unknown functions as the most diverse within the SfMNPV genome. Furthermore, an evolutionary study of *sf29* suggests that this gene encodes a putative collagenase with a conserved catalytic motif and presents evidence of pervasive positive selection. We detected several structural variants (mostly deletions and duplications) that contribute to the genomic diversity of SfMNPV. Comparative genomic analysis based on a whole-genome alignment placed ARG-M close to Nicaraguan isolates and suggests that single nucleotide variants (SNPs) are useful to discriminate the genomic composition of different isolates, as demonstrated previously for CpGV (Alletti et al. 2017). We believe that our data will help to better understand the adaptation process of SfMNPV to the fall armyworm and would be useful to improve the applicability of SfMNPV as biopesticide.

## 2. Materials and Methods

### 2.1. Virus isolation and DNA sequencing

The argentine isolate SfMNPV (ARG-M) was isolated from a single S. frugiperda larva collected from a corn field in Oliveros (Santa Fe Province) in 1985 and further purified at the IMyZA, INTA, Castellar, Argentina (Berretta, Rios, and Cap 1998). For virus amplification, a laboratory *S. frugiperda* colony was established at the IBBM, La Plata. Virus stock was amplified in 3 day old larvae fed with virus contaminated artificial diet and maintained individually at 26°C, 12:12 h (light:dark) photoperiod until death or pupation. Dead larvae were homogenized in double-distilled water (ddH_2_O), gauze filtered and resuspended in 0,1% sodium dodecyl sulfate. Cell debris was eliminated by low speed centrifugation (1000g, 2 min) and supernatant with viral occlusion bodies (OBs) was centrifuged at 12000 g. OBs were washed twice with ddH_2_O and resuspended in ddH_2_O. OBs were loaded on to a discontinuous 35–60% (w/w) sucrose gradient and centrifuged at 20000 rpm and 4°C for 1 hour in a SW41 rotor (Beckman). The band containing the OBs was removed, washed twice in ddH_2_O and the viral pellet was resuspended in ddH_2_O. Purified OBs were dissolved by alkaline lysis and DNA extraction was performed as described previously (Ferrelli et al. 2018). SfMNPV-ARG M genomic DNA was used for library preparation (TruSeq DNA PCR-Free kit) and sequenced on a Illumina HiSeq 4000 platform at Macrogen Corporation (South Korea). Raw sequencing dataset corresponds to 70239480 reads, which are deposited at the NCBI SRA database under accession number PRJNA648791.

### 2.2. Genome assembly and annotation

Genome assembly pipeline was based on the Snakemake workflow manager in order to ensure a consistent and reproducible analysis (Köster and Rahmann 2018). Quality checking and adapter trimming from raw reads was carried out simultaneously with fastp, using default parameters (Chen et al. 2018). In order to speed-up the genome assembly, we sampled 2 million reads from the original dataset for the subsequent steps using the seqtk tool. Genome assembly was performed with Megahit (Li et al. 2015) and the longest scaffold was selected for further refinement. Custom scripts were used to circularize the scaffold and set the polyhedrin start codon as the genome start position. Putative open reading frames (ORFs) were predicted with ORFfinder with the following parameters: -c t -s 0 -ml 150 (Sayers et al. 2010). Coding sequences annotation was carried out using the InterProScan tool (Jones et al. 2014) and BLASTp homology search (Camacho et al. 2009) against available SfMNPV genome sequences followed by annotation data retrieval using the Entrez Direct utilities (Sayers et al. 2010). Genome sequence data was deposited at GenBank under accession number MW162628.

### 2.3. Single nucleotide and structural variants calling

All reads from the complete dataset were processed with fastp (Chen et al. 2018). Reads that passed the quality filter were aligned against the viral genome assembly using the BWA-MEM algorithm (H. Li and Durbin 2009) and subsequently sorted with SAMtools (H. Li et al. 2009). Lofreq2 was used for single nucleotide variant calling and variant effect was evaluated using SnpEff together with our previous annotation as reference (Wilm et al. 2012; Cingolani et al. 2012). Delly (Rausch et al. 2012) and Lumpy (Layer et al. 2014) were invoked with default parameters in order to detect evidence of structural variation present in our mapped reads dataset.

### 2.4. Phylogenetic analysis

Genome and proteome sequences from all available SfMNPV isolates were downloaded from NCBI nucleotide and protein databases, respectively. A whole-genome alignment was constructed with MAFFT (Katoh and Standley 2013) and SNPs were extracted from the genome alignment with SNP-sites (Page et al. 2016). A maximum likelihood phylogeny was inferred with IQ-TREE with default settings. Model selection was done with ModelFinder and 1000 replicates of ultrafast bootstrap were used to calculate branch support (Minh et al. 2020; Kalyaanamoorthy et al. 2017; Hoang et al. 2017). Tree manipulation, coloring and visualization was done with ToyTree (Eaton 2019) and Inkscape. Protein sequences were clustered using BLASTp (Camacho et al. 2009), aligned with MAFFT (Katoh and Standley 2013) and then used to extract nonsynonymous mutations affecting coding sequences.

### 2.5. Molecular evolution analysis of *sf29*

BLASTp homology search (Camacho et al. 2009) was used to identify orthologs of *sf29* present in baculovirus and entomopoxvirus. Bacterial orthologous sequences were included as outgroups for the phylogenetic analysis. Protein sequences were retrieved for each ortholog and used to construct a sequence alignment and a phylogenetic tree, as described above. Molecular evolution analysis was carried out with codon-level alignments using the HyPhy package (Pond et al. 2019). We checked for the absence of recombination in our alignment with GARD (Pond et al. 2006). To detect signatures of pervasive and episodic positive selection for each site in the codon alignment we used FEL (Pond and Frost 2005) and MEME (Murrell et al. 2012), respectively. Evidence of diversifying selection in the different branches of our phylogenetic tree was studied with aBSREL (Smith et al. 2015). Because some of the evolutionary analysis mentioned before are computationally intensive, we used the Datamonkey web-server (Weaver et al. 2018) to run some of them. Sequence disorder prediction was performed with IUPRED2 (Mészáros, Erdős, and Dosztányi 2018) and alignments visualization with Aliview (Larsson 2014).

### 2.6. Code and data availability

Reads and genome sequence data have been deposited at NCBI SRA and GenBank databases, respectively. Code used to process data and generate figures is available at Github (tomasMas-son/sfmnpv_genomics). Output files generated with HyPhy can be visualized interactively using the web tool HyPhy Vision.

## 3. Results

### 3.1. Deep sequencing of SfMNPV ARG-M

An argentinian autochthonous isolate of SfMNPV, named ARG-M, was characterized through deep sequencing using Illumina technology for the first time, enabling the study of its genomic diversity. Our deep sequencing data reached an average coverage of 52313X (standard deviation of 4052). The consensus genome sequence of SfMNPV ARG-M is 132696 base pairs (bp) long and comprises 144 coding regions (Accession number MW162628, Supplementary Table 1), from which 143 are present in the reference isolate 3AP2 (Harrison, Puttler, and Popham 2008) and the remaining gene corresponds to the ORF *sf110a*, present only in Nicaraguan isolates (NicB and NicG) (Simón et al. 2011). When genome structural differences between ARG-M and the reference isolate were examined, we only found a small number of insertions/deletions shorter than 6 bp at non coding regions, with the exception of a 31 bp deletion at the intergenic region between *me53* and *hr1*. The remaining discrepancies between both genome sequences were due to SNPs.

### 3.2. Genome-wide diversity in a SfMNPV isolate

From our sequencing data we could detect 704 intrahost single nucleotide variants (iSNVs) and their associated allele frequencies, as reported by LoFreq (Supplementary Table 2). Global distribution of iSNVs was homogeneous across the viral genome and encompassed a wide range of allele frequencies (Fig. 1a). Genetic diversity of our SfMNPV population was 5.3 × 10^−3^ iSNVs per bp. We classified iSNVs into three groups based on its predicted effect on coding sequences (nonsynonymous, synonymous and intergenic). Approximately half of these iSNVs correspond to synonymous variants (384), while the remaining are distributed between nonsynonymous (184) and intergenic variants (131) (Fig. 1b). Additionally, we found four variants that introduced premature stop codons (Glu357* *sf41*, Glu392* *sf43*, Tyr73* *sf47* and Cys6* *sf110a*) and one variant which abolished a start codon (*sf45*). As expected, synonymous variants presented a higher median frequency of the three variants groups, probably because of their mild impact in comparison with nonsynonymous mutations affecting coding sequences or variants affecting regulatory regions (Fig. 1c). Remarkably, 31% (54 of 184) of nonsynonymous iSNVs showed an allele frequency greater than 0.25, which suggests that significant levels of different proteoforms of the same gene can coexist during SfMNPV infection.

**Figure 1:**
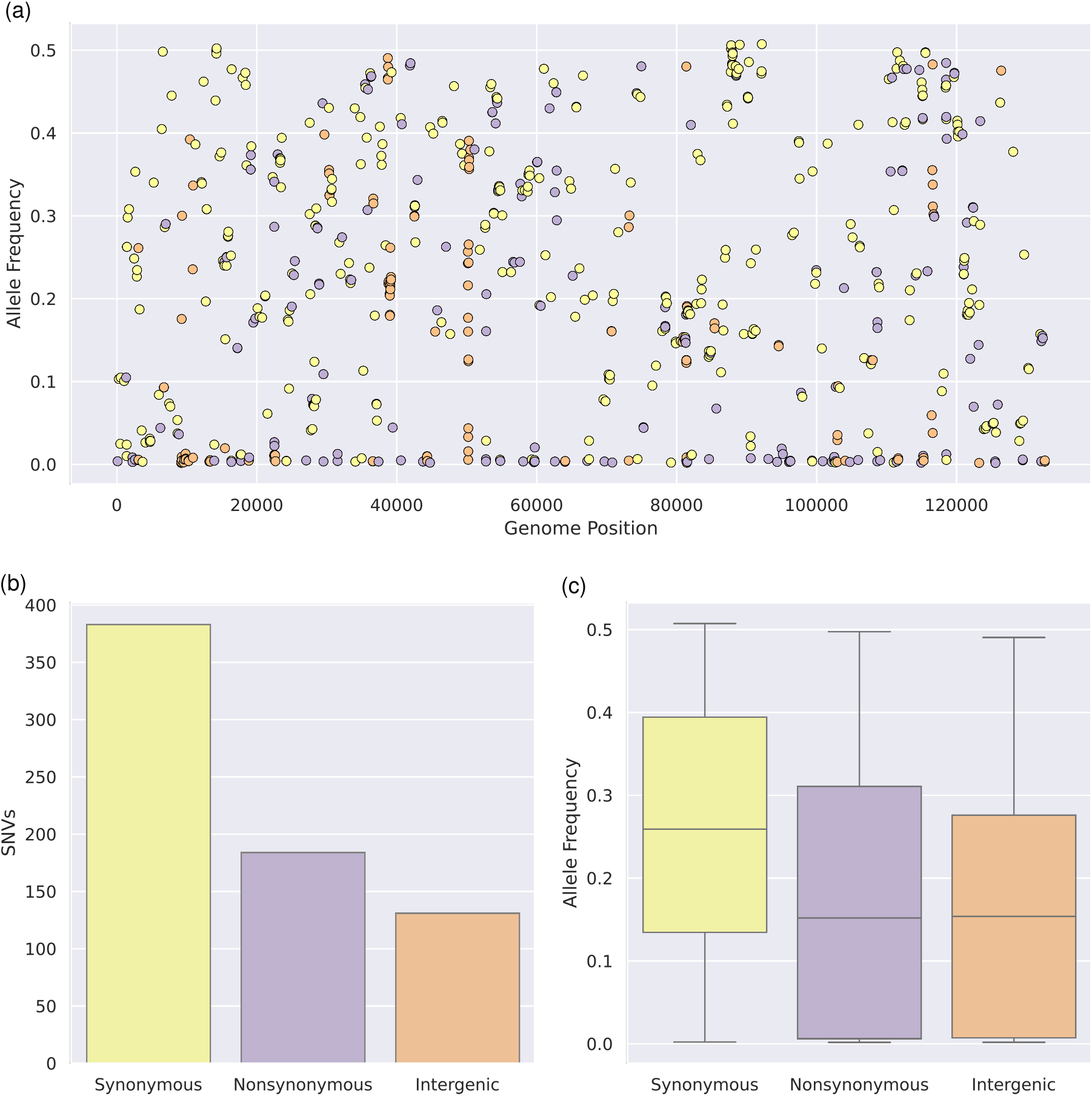
Genome-wide diversity in the SfMNPV ARG-M isolate. **(a)** Distribution of synonymous (yellow), nonsynonymous (violet) and intergenic (orange) iSNVs across the genome. **(b)** Number of iSNVs detected for each category. **(c)** Boxplot showing the allele frequency for each iSNV category.

Based on the number of nonsynonymous iSNVs, SfMNPV genes were ranked according to their genetic diversity. We identified *sf29*, *odv-e66a*, *odv-e66b*, *sf23*, *p40*, *pif-0*, *sf110a*, *lef-6*, *lef-7*, *sf68*, *vp80*, *rr1*, *94k* and *p47* as the 10% most variable genes. Another observation that emerges from our data is that approximately 57% (105 of 184) of nonsynonymous iSNVs introduced a change in amino acid polarity, which could lead to modifications in protein properties, as described previously (Chateigner et al. 2015). From these polarity-changing mutations, only 27 affected core genes while 78 were located inside non-core genes, highlighting the greater restriction of core genes to incorporate drastic changes on its sequence.

Genome structural variants (SVs), which comprise deletions, insertions, inversions, duplications and translocations, represent another source of diversity in populations of baculovirus and other large DNA viruses (Loiseau et al. 2020). For SfMNPV, deletions affecting the *egt* locus have been reported previously as the main SVs (Serrano et al. 2012; Niz et al. 2020). We decided to investigate the presence of other SVs in our dataset in order to complement the existing body of knowledge. Using Delly and Lumpy as SV callers, we retained 13 SVs detected with single-base precision by both callers (Supplementary Table 3). We recovered two large deletions comprising the region 22000-26000 that erased completely the *egt* gene, in agreement with previous reports. In addition, novel SVs comprised two large deletions that ablated significant portions of the genome (36.5 and 54.2 Kbp), five small deletions (affecting *f-protein*, *pif-1*, *odv-e66a* and *cg30*), one inversion of 52.8 Kbp and three duplications (4.3, 5.0 and 9.8 Kbp). Using read support for each SV and the mean coverage of our dataset, we estimated the frequency per genome for each SV, as described elsewhere (Gilbert et al. 2014). Duplications and most of the deletions occurred with a frequency in the range 1.9 × 10^−5^-1.7 × 10^−3^, but deletions affecting *egt* and *odv-e66a*, together with the inversion event, presented higher frequencies (2.3 × 10^−3^-1.2 × 10^−2^).

### 3.3. Genetic diversity of SfMNPV isolates

Five genome sequences from SfMNPV isolates are available to date. Genomes B and G of the Nicaraguan isolate have some distinctive characteristics regarding their gene content compared to 3AP2 and 19 genomes (Simón et al. 2011). So, we compared these features in ARG-M. There are three ORFs reported only in NicB and NicG, that are absent in 3AP2 and 19, namely *sf39a*, *sf57a* and *sf110a*. Of these, we found *sf110a* in ARG-M but not *sf39a* or *sf57a*. *Sf39a* is a 63 aa protein in NicB and NicG. TBlastn search revealed that ARG-M has a shorter homolog of 33 aa, and therefore it was not annotated. *Sf57a* is located in an insertion reported in NicB, which is absent in 3AP2 and 19 (Simón et al. 2011), as well as in ARG-M. On the other hand, ARG-M codes for *sf129*, a gene that was exclusive of 3AP2 and is absent in Nicaraguan genomes as well as in 19. Therefore, ARG-M presents a different gene pattern regarding these differential genes in comparison to the other SfMNPV genomes.

We constructed a whole-genome multiple sequence alignment (MSA) in order to explore the underlying genomic diversity between different isolates. Phylogenetic reconstruction resulting from this MSA showed that SfMNPV isolates are highly similar, with evolutionary distances in the order of 1 × 10^−3^ substitutions per bp. In this context, the ARG-M isolate is more similar to the Nicaraguan isolates, whereas the isolates 19 and 3AP2 cluster together and the Colombian isolate represents the most divergent SfMNPV genome (Fig. 2a). Based on the methodology reported by Wennmann et al. (2017), where polymorphism data was used to cluster different *Cydia pomonella* granulovirus (CpGV) isolates, we extracted 1502 SNPs present in our MSA (Supplementary Table 4) and analyzed its distribution across the SfMNPV genome. We evidence an homogeneous distribution with an approximate of 100 SNPs per 10.34 Kbp bin, except for the region 20000-30000 where a peak of SNPs occurred (Fig. 2b). Upon further inspection, this higher number of SNPs could be explained by the highly divergent genomic segment of *Spodoptera litura* II nucleopolyhedrovirus (SpltNPV-II), introduced by an horizontal gene transfer event into the Colombian isolate (Barrera et al. 2015). Also, we noticed an appreciable number of isolate-specific SNP: Colombian (577), 19 (210), 3AP2 (162) and ARG-M (87) (Fig. 2c). Although we found 93 SNPs common to isolates NicB and NicG, we couldn’t identify SNPs specific to NicB and NicG displayed only 16 unique SNPs. In general, we observed that SNPs shared by two or three isolates are in agreement with our phylogenetic reconstruction: isolates ARG-M, NicB and NicG share 92 SNPs, while isolates 19+COL and 19+3AP2 have in common 59 and 56 SNPs, respectively.

**Figure 2:**
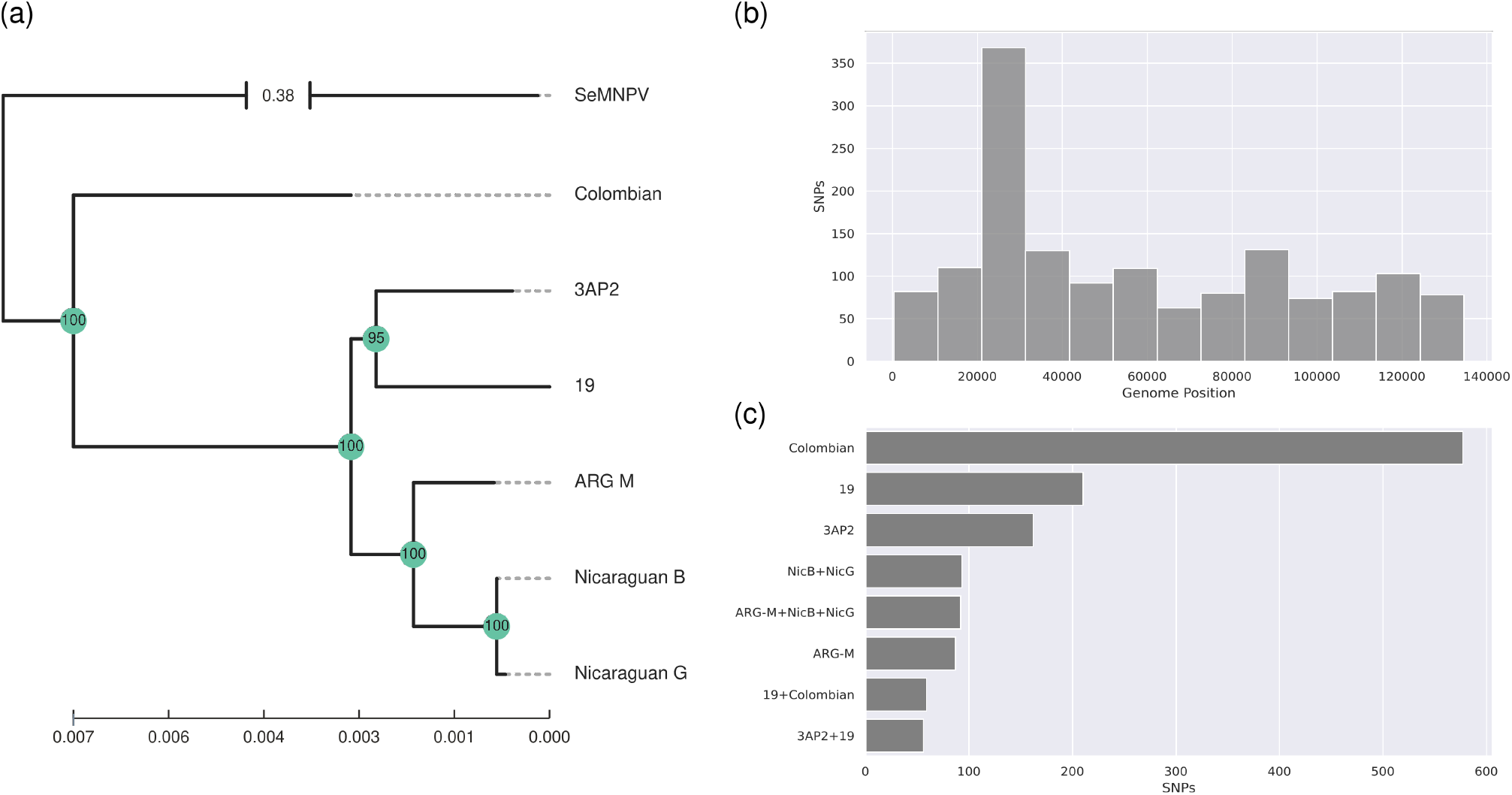
Genetic diversity present in SfMNPV isolates. **(a)** Maximum likelihood phylogeny reconstructed from a whole-genome sequence alignment. Support values were computed using 1000 Ultra-Fast Bootstrap replicates. *Spodoptera exigua* nucleopolyhedrovirus (SeMNPV) was selected as outgroup. **(b)** Histogram showing the number of SNPs per bin detected within the sequence alignment. The alignment was splitted into 13 bins, which correspond to 10.38 Kbp per bin. **(c)** Barplot showing the number of SNPs detected for each isolate and the combinations detected.

Because we were interested in unraveling which coding sequences have the largest variability, we analyzed the diversity levels of inter- and intraisolate nonsynonymous mutations for each coding region (Fig. 3). In general, we noticed that iSNVs and SNPs tend to affect the same positions within coding regions. Most genes showed low levels of variation, with less than 3 × 10^−3^ nonsynonymous mutations per bp. However, 15 genes presented diversity levels above 3 × 10^−3^ nonsynonymous mutations per bp for inter- and intraisolate datasets (*lef-7*, *sf23*, *sf29*, *odv-e66a*, *p40*, *sf68*, *gp41*, *sf85*, *sf110a*, *sf118*, *sf122, ubiquitin*, *sf126*, *lef-6* and *sf135*). Several of these genes are coincident with our diversity rank based solely on nonsynonymous iSNV counts.

**Figure 3:**
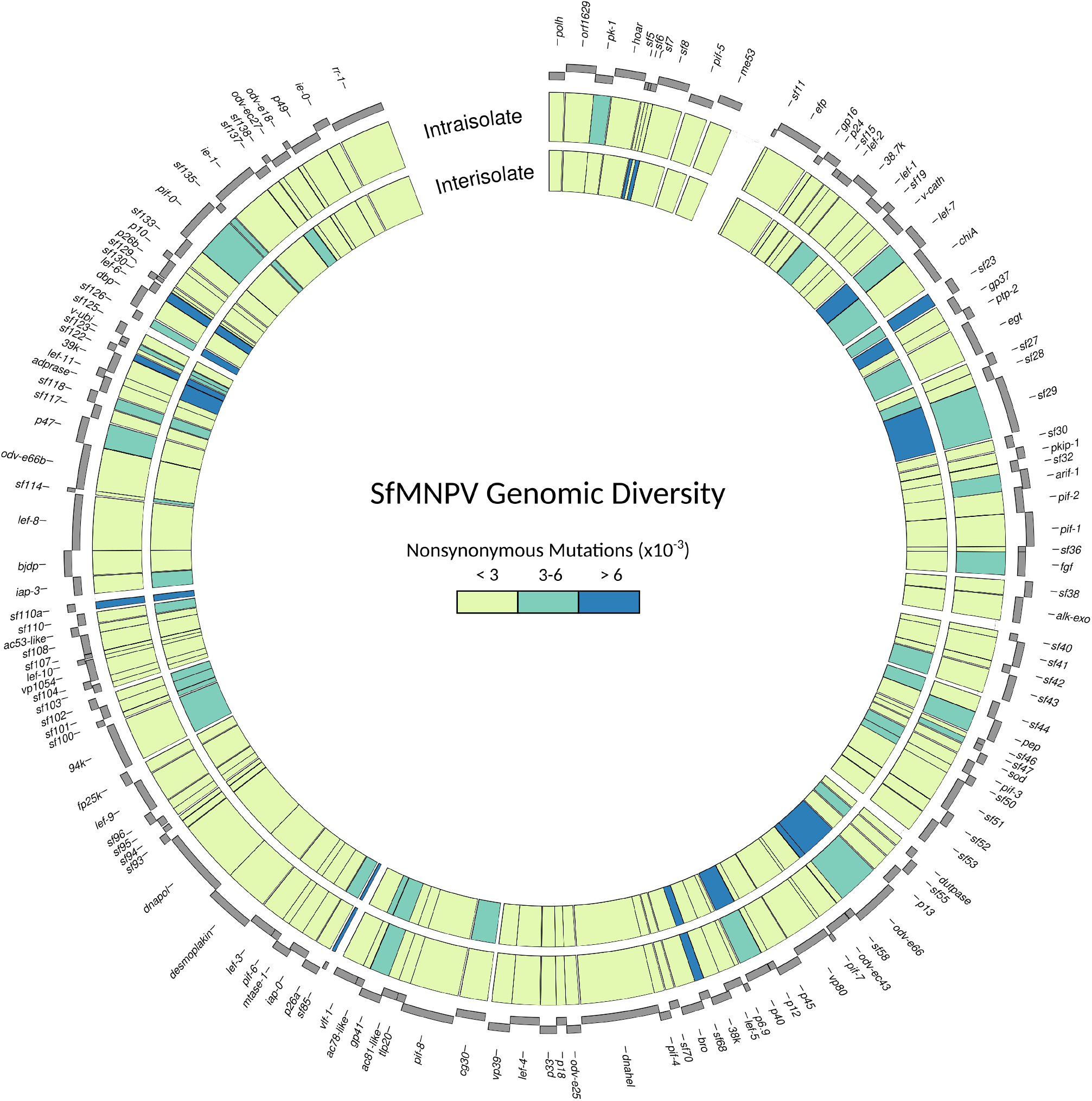
Proteome diversity within SfMNPV. For each coding region in SfMNPV the number of nonsynonymous variants was computed. The inner circle represents SNPs derived from isolates whole-genome sequence alignment, while the outer ring corresponds to the iSNVs detected within SfMNPV ARG-M.

### 3.4. Molecular evolution of *sf29* in insect virus

Based on our previous results, we decided to characterize the evolutionary process of *sf29* in order to evidence signals of adaptive evolution. This protein has been described as a viral factor determining the number of virions present inside OB particles and it is distributed in group II *Alphabaculovirus* II and *Entomopoxvirus* (Simoń et al. 2008). A maximum likelihood phylogeny of *sf29* suggests a possible bacterial origin for this protein and a subsequent transfer event to an ancestral insect virus before passing to baculovirus and entomopoxvirus (Fig. 4a). Furthermore, the aBSREL method found evidence of diversifying selection acting specifically on several *Entomopoxvirus* branches (MySEV, AHEV and two AMEV species, highlighted in orange), which suggests a sustained regime of diversifying selection within this clade. The baculovirus branches corresponding to PespNPV and MyunNPV also showed signs of diversifying selection.

**Figure 4:**
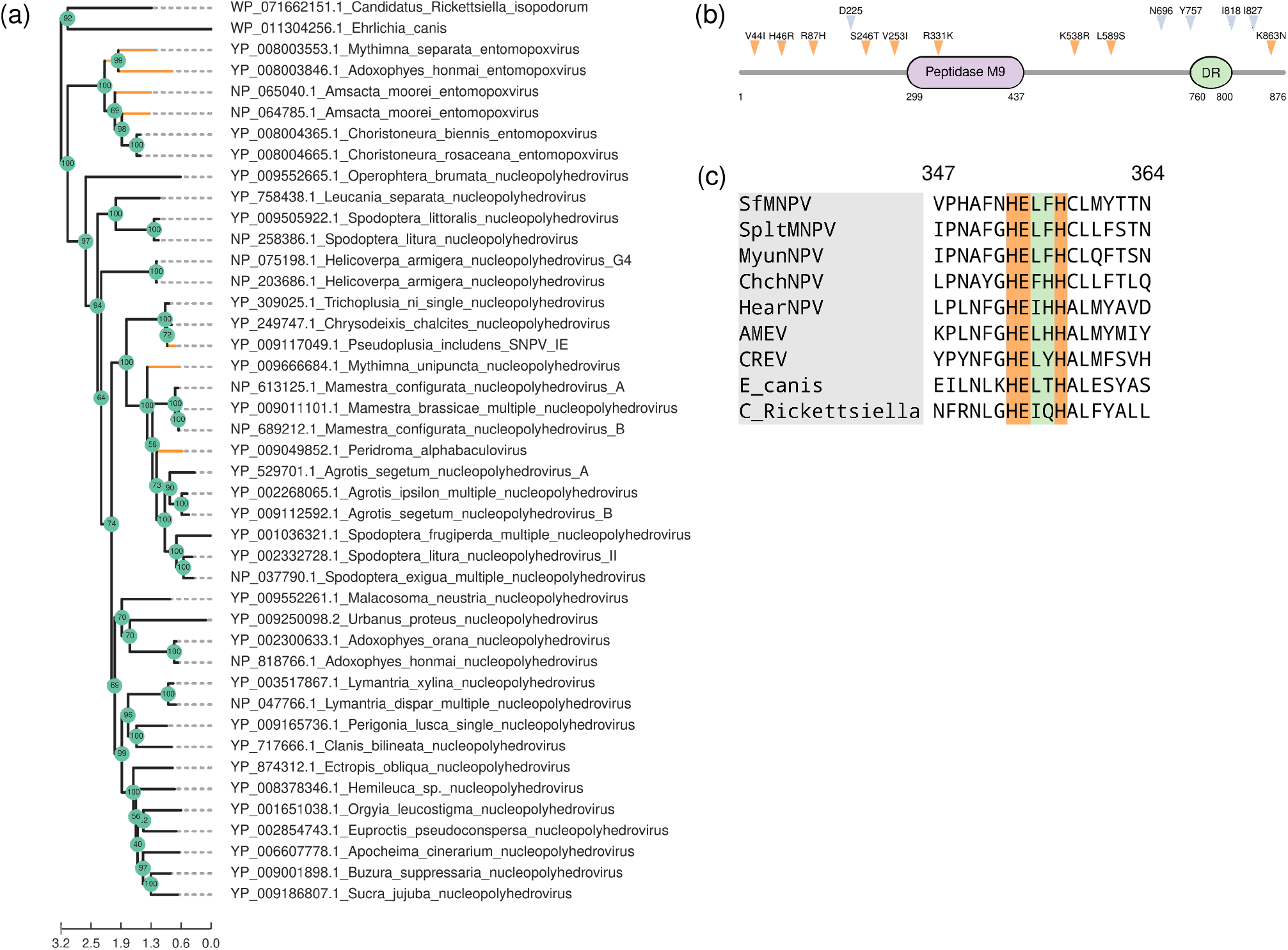
Sf29 Molecular Evolution. **(a)** Maximum likelihood phylogeny of *sf29* orthologs. Support values were computed using 1000 UltraFast Bootstrap replicates. *E. canis* and *C. Rickettsiella* were included as outgroups. Branches experiencing positive selection, as detected by aBSREL, are colored in orange. **(b)** Sequence analysis of *sf29*. Sites under positive selection are marked in blue, while iSNV are depicted in orange. The peptidase M9 domain is colored violet and the disordered region is showed in green. **(c)** Sequence conservation of the peptidase activate site motif (HEXXH), extracted from the *sf29* sequence alignment.

Sequence analysis revealed the presence of a Peptidase M9 (collagenase) domain (residues 299-437) and a disordered region (760-800) in *sf29* (Fig. 4b). Additionally, a signal peptide and a C-terminal transmembrane region were detected, suggesting that *sf29* could be a membrane protein. Positive selection test performed with FEL showed evidence of positive selection for five residues (D225, N696, Y757, I818 and I827;). These positively selected residues and the iSNVs detected in the ARG-M population are located mainly outside the predicted peptidase domain, which suggest that adaptive evolution in *sf29* is driven by its non-peptidase regions. Moreover, several residues experiencing episodic positive selection, as tested by MEME, were coincident with iSNVs present in the SfMNPV ARG-M population. Analysis of residue conservation revealed that the HEXXH motif, involved in zinc binding and a common feature of metalloproteases (Rawlings et al. 2013), is conserved in most of the sequences of our dataset, suggesting that *sf29* could be a functional collagenase (Fig. 4c).

## 4. Discussion

It is increasingly recognized the relevance of genomic diversity as a key determinant of viral evolutionary dynamics and virulence (Geoghegan and Holmes 2018), specially in large DNA viruses (Renner and Szpara 2017). This work presents a snapshot of the underlying genomic diversity present inside a natural isolate of SfMNPV, expanding previous studies of isolates for which only a consensus sequence has been reported (Harrison, Puttler, and Popham 2008; Wolff et al. 2008; Simón et al. 2011; Simón, Palma, et al. 2012; Barrera et al. 2015).

The genetic diversity found in SfMNPV ARG-M (5.3 × 10^−3^ iSNV per bp) is similar to previous reports of other baculovirus species (Brito et al. 2015, 2018; Alletti et al. 2017; Zwart et al. 2019). Because non-synonymous variants have the potential to introduce drastic changes on protein structure, we focused much of our analyses on this type of mutation. Both iSNVs and SNPs pinpointed a common subset of genes (Fig. 3) with high levels of genetic diversity. Most of these genes (*sf23*, *sf85*, *sf110a*, *sf118*, *sf122*, *sf126*, *sf135*) encode proteins with unknown function, indicating that a great source of adaptability remains uncharacterized in SfMNPV. Examination of these genes with the InterProScan tool only found a disordered region within *sf23* (residues 180-209) and predicted a domain of unknown function (DUF424, e-value 9.5 × 10^−20^) inside *sf126*. In addition, three genes associated with oral infectivity (*odv-e66a*, *sf68* and probably *sf29*) also presented an elevated genetic diversity. *Odv-e66a* encodes a chondroitin lyase that degrades peritrophic membrane, while *sf68*-derived protein contains a chitin-binding domain (Superfamily SSF57625, e-value 5.49 × 10^−5^) and *sf29* carries a collagenase domain. In this scenario, it is possible that the elevated genetic diversity of these viral genes could be useful to maintain an optimal adaptation to the hostile environment imposed by the midgut of the insect host (Daugherty and Malik 2012; Wang, Zhao, and Han 2020).

Deletions located in the *egt* locus are common in natural populations of SfMNPV and have been related with the speed of kill and the potency of occlusion bodies inoculum (Simón, Williams, et al. 2012). However, there is little information addressing the genome-wide occurrence of SV on SfMNPV. Here, we found several SV comprising deletions, duplications and one inversion. One of the most frequent SV detected was the deletion affecting the *egt* locus, an observation that could be related with the sustained detection of this deletion in natural populations. Another point that arises from our reported SV is that deletions were located roughly within one half of SfMNPV genome sequence (position 6505 to 73275), while duplication events affected genome positions on the other half of genome sequence (positions 80710 to 121444). Inspection of the gene content for each of these two regions evidenced that deletions conduct to the loss of genes associated mostly with oral infectivity, auxiliary functions or unknown functions, while duplications results in the increase of genes related with replicative/transcriptional functions (*ie-1*, *dnapol*, *lef-3* and *lef-8*). This differential localization of deletions and duplications supports a hypothetical picture where virus multiplication could be accelerated by a subpopulation of viral genomes with reduced size and higher copy number of replicative/transcriptional genes, as proposed previously (Rezelj, Levi, and Vignuzzi 2018; Leeks et al. 2018).

Motivated by the presence of a putative collagenase domain inside *sf29* and its elevated genetic diversity, we conducted a detailed evolutionary study of this gene to better understand its possible function. *Sf29* evolutionary history exemplifies the recurrent gene flow between baculovirus and other organisms (Thézé et al. 2015; Rodrigues et al. 2020), in this case, with *Entomopoxvirus* and probably prokaryotes. In addition to its predicted collagenase domain, we demonstrated the pervasive conservation of the HEXXH motif, a hallmark of metalloproteases (Rawlings et al. 2013), which reinforces the hypothesis that *sf29* encodes a functional collagenase. Through different methods we evidenced the presence of positive selection acting on *sf29*. One possible explanation for the adaptive evolution detected in this gene could be related with the different selective environment experienced by *sf29* after its capture by an ancestral insect virus, a process that is more pronounced in *Entomopoxvirus*, according to our aBSREL results. Further molecular studies of *sf29* are necessary to confirm its biochemical activity and its role during oral infection.

Polymorphism exhibited by SfMNPV isolates represents a useful tool to catalogue different isolates and to ensure their field efficacy. Isolate genotyping is an important strategy to manage the emergence of isolate-dependent resistant populations of the codling moth, as demonstrated by the continued use of CpGV (Alletti et al. 2017). Although the number of SfMNPV isolates is still low, some generalities can be extracted from its SNPs pattern. We found that most SNPs present in SfMNPV isolates are biallelic and specific to only one isolate, suggesting that SNPs could be used for genotyping novel isolates. Recently, genetic variation between nudivirus isolates with different viral titers have been used to select SNPs that are directly linked to virulence (Hill and Unckless 2020), a result that highlights the usefulness of sequencing data to infer biological properties. With the increasing access to HTS infrastructure and the availability of computational tools to analyze the data generated, we believe that sequencing of SfMNPV isolates across geographical and temporal scales will improve our knowledge of the molecular determinants of SfMNPV pathogenicity. This work provides the first study addressing the underlying genomic diversity in a natural isolate of SfMNPV and its possible consequences on virus adaptation to the fall armyworm.

## Supplementary Tables

**Supplementary Table 1:** Annotated coding sequences for SfMNPV ARG-M.

**Supplementary Table 2:** Intrahost single nucleotide variants (iSNVs) detected with Lofreq.

**Supplementary Table 3:** Structural variants detected with both Delly and Lumpy.

**Supplementary Table 4:** Single nucleotide polymorphisms (SNPs) present in SfMNPV isolates.

